# Genetically informed distribution models refine predictions for the overwintering range of *Helicoverpa armigera* in North America

**DOI:** 10.64898/2026.08.21.746245

**Authors:** Cian D. Williams, Chris D. Jiggins, Henry L. North

## Abstract

The ecological and economic threat posed by invasive pests demands proactive mitigation. Species distribution models (SDMs) are widely used in efforts to predict where invasive species might spread after introduction, though such models face several limitations. Among these is the unrealistic assumption of niche uniformity throughout a species’ range. This has led to interest in developing SDMs that explicitly account for local adaptation, though few methods have achieved this in a way that confidently separates local adaptation from population structure. Here we develop and implement a sequential SDM approach that incorporates experimentally verified associations between genotype, phenotype, and environment to forecast establishment risk in a major agricultural pest. We leverage genomic data from 738 individuals to characterize the geographic distribution of alleles at a major-effect locus for cold tolerance *(tret1)* in *Helicoverpa armigera*, an invasive crop pest of major economic concern in North America. We demonstrate that a recently detected North American population carries a cold-adapted *tret1* allele, which has likely contributed to its persistence. We quantify the contribution of cold-adapted *tret1* to the potential invasive range of *H. armigera* in North America under current and future climate scenarios. We find that cold-adapted *tret1* may dramatically expand the potential range of *H. armigera*, and that potential future range expansion is likely to be driven primarily by cold-adapted individuals. Our results highlight the importance of accounting for intraspecific variation in invasive species risk assessments and management strategies.

**Significance Statement:** *Helicoverpa armigera* is an agricultural pest of global economic concern. The species is invasive in South America, and is likely to spread to North America, where crops worth an estimated US$843 million per annum are grown in regions considered to have an optimal climate for the pest (1). However, predictions to date struggle to account for the fact that *H. armigera* populations are locally adapted to different climatic conditions. Leveraging a large genomic dataset, we demonstrate that the first known population in North America carries an allele conferring increased cold tolerance. We use sequential distribution modeling — an approach that allows the integration of diverse data sources to predict habitat suitability — to incorporate this information, refining forecasts of its potential range in current and future climate scenarios. We find that pre-existing cold tolerance likely contributed to the persistence of the small, localized resident *H. armigera* population in North America, and could facilitate its spread of throughout much of the agriculturally productive American Midwest.

## Introduction

Invasive species pose one of the most significant threats to biodiversity and ecosystem services globally (2). Assisted by human travel and trade in the Anthropocene (3, 4), the spread of non-native species continues to accelerate. In addition to the extensive damage they cause to native biodiversity (5, 6), the spread of invasive species can also result in large economic burdens in agriculture (7), fisheries (8), and public utilities (9), with global costs estimated at over $400 billion USD annually (10).

Management responses to the detection of an invasive species in an area outside its known range often include an assessment of habitat suitability (11). Such analyses inform management decisions: if the habitat in a putative non-native range is deemed unsuitable for the species, then little intervention may be needed; otherwise swift action is required to prevent the species establishing and spreading further (12). Habitat suitability predictions are also useful prior to detection, enabling targeted resource prioritization planning. Species distribution models (SDMs) are a class of machine learning methods often used for this purpose (13). In the context of risk assessments for invasive species, SDMs are often fitted using presence-only data in a species’ native range to identify environmental predictors of habitat suitability, before being used to predict habitat suitability in the potential non-native range.

A key limitation of SDMs is their assumption of niche uniformity throughout a species’ range, with no local adaptation (14). This is particularly problematic for modeling distributions of invasive species, where the introduced population is often founded by a small number of individuals, likely from the same non-native region and with limited genetic variation (15). This means that any locally adaptive traits in the source population may have a large impact on the probability, rate, and extent of spread in the non-native range. As high-throughput DNA sequencing has become cheaper and more accessible, there has been growing interest in developing SDMs that incorporate information about local adaptation from genotypic data (16, 17). However, the substantial additional effort required to sample genotypic data can lead to limited geographic sampling, which can in turn lead to truncation of the inferred niche in an SDM, resulting in misleading predictions (18).

Several methods have recently been developed for incorporating local adaptation into predictions of species ranges and responses to climate change. Arguably the most prominent example is the growing family of “genomic offset” methods. These methods identify putatively adaptive loci as outliers in gene-environment association tests, then use these to predict the hypothetical combination of observed alleles that would maximize fitness in a new habitat or under future climatic conditions (19). The resulting genomic offset — the difference between the current and putatively optimal genetic composition — is used as a proxy for maladaptation. An advantage of these approaches is their use of genome-wide data. However, even if gene-environment association can perfectly distinguish between locally adapted genetic variation and non-adaptive population structure, genomic offsets suffer from difficult interpretability issues, as the link between genomic offset and fitness or population vulnerability is unclear (20). Other attempts to incorporate local adaptation have included identifying genetic clusters within species and modeling them separately in an SDM (21–23). However, these approaches frequently do not differentiate between spatial genetic structure arising from local adaptation and that arising from isolation by distance or other non-adaptive processes. Further, both approaches are still limited by the small sample sizes necessitated by population genomic studies, making them vulnerable to niche truncation. An ideal approach would consider the distribution of individuals carrying a locally adapted genotype, while also incorporating information about the global distribution of the species, to mitigate niche truncation. We propose that spatially nested SDMs (N-SDMs), which have previously been used to integrate broad-scale species occurrences with local-scale occurrences to overcome niche truncation (24), are well-suited for this problem. Here we use a sequential modeling approach to produce a genotypically informed N-SDM for an agriculturally significant invasive crop pest that accounts for local adaptation.

The Old World bollworm *Helicoverpa armigera* (Lepidoptera: Noctuidae) is one of the world’s most economically devastating agricultural pests, causing an estimated $4billion USD of crop damage annually (19). It is a highly polyphagous pest of economically important crops including soy, cotton, and maize (20) and has evolved resistance to many commonly used pesticides (19). In its native range, *H. armigera* is broadly distributed across Africa, Eurasia, and Australia (21), but was declared an invasive species throughout much of South America in 2013 (22). In 2019, an individual specimen of *H. armigera* was captured during a routine port environs survey at Chicago O’Hare International Airport (ORD) in the United States (23), apparently because of an escape from air freight. In each subsequent year, additional specimens were captured in the area immediately surrounding ORD, indicating the presence of a small, geographically restricted, reproducing population. This population retained high genetic diversity and appears to have been introduced from a source population in South America, though the possibility of multiple introductions to ORD has not been ruled out (24). The detection of this population has led to renewed interest in reassessing the risk of spread throughout North America, especially given that the genotype of the founding population may influence its potential range — a consideration overlooked in previous range forecasts for this species.

The generalist ecology of *H. armigera* combined with a tendency for long-distance dispersal means that there is remarkably low genetic differentiation across its vast native range (25). Therefore, the possibility of substantial local adaptation in this species appeared minimal. However, recent phylogeographic studies incorporating global geographic sampling of whole-genome resequencing data challenged this finding (26, 27). Jin *et al*. (26)identified an autosomal locus including the trehalose regulator *tret1* at which allelic variation was strongly associated with minimum winter temperatures, driven largely by alleles at high frequency in the Xinjiang region of north-western China where winter temperatures fall below −20°C. Trehalose is the main hemolymph sugar in insects (28) and also acts as a compatible solute, inhibiting the formation of ice crystals at low temperatures. The same study provided three additional lines of evidence for local adaptation at the *tret1* locus. First, allele frequencies are highly differentiated at this locus between samples collected in Xinjiang and nearby populations in warmer regions despite little differentiation across the rest of the genome. Second, a selective sweep was detected at this locus in the Xinjiang population. Third, pupae carrying the cold-adapted genotype transport trehalose more efficiently, enabling them to maintain higher concentrations of trehalose and withstand colder temperatures before the formation of ice crystals when compared to carriers of the wild-type (WT) genotype on the same genetic background. Therefore, *H. armigera* homozygous for the cold-adapted *tret1* allele transport trehalose more efficiently, enabling them to withstand colder temperatures through cryobiosis, which increases fitness in regions with very low winter temperatures. The cold-adapted and wild-type *tret1* alleles differ by only a single nonsynonymous SNP (1190G>A). Below we use ‘WT’ and ‘cold adapted’ to refer to the two genotypes.

Given the low winter temperatures in Chicago, Illinois, where the North American *H. armigera* population is currently restricted, genetic variation at *tret1* in this population has clear implications for its probability of persistence and the geographic extent of its potential overwintering range in North America. Here, we take advantage of the availability of whole-genome resequencing data and knowledge of locally adapted alleles in *H. armigera* to explore the use of a sequential modelling approach, producing a genotypically informed N-SDM models of this economically significant pest. We demonstrate that the introduced population carries the cold-adapted *tret1* allele at high frequency, likely due to selection at ORD. Using a sequential SDM approach, we quantify the contribution of cold-adapted *tret1* to the potential range of *H. armigera* in North America, both in the present and under projected future climate scenarios.

## Results

### Spatial sorting of cold-adapted *tret1* alleles in native and invasive populations

Using whole-genome resequencing data from *H. armigera* collected throughout its native and introduced ranges (N = 738), we called variants at the *tret1* locus to characterize the geographic distribution of the cold-adapted *tret1* allele (Fig. 1A). In the native range, cold-adapted *tret1* is present at high frequency (allele frequency=92.6%, N=82 individuals) in the Xinjiang region of northwest China, which experiences winter temperatures below −20°C (26). There is a geographic cline in cold-adapted allele frequency from northern to southern China, with the cold-adapted allele present at low frequency in southern China (12.0%, N=50) despite low genetic differentiation between these populations (26). The cold-adapted allele is present at low (<10%) frequency throughout the rest of the sampled native range in Australasia, south Asia, Africa and southern Europe. The persistence of this cline in the face of gene flow confirms multiple previously established lines of evidence for local adaptation at this locus.

**Figure 1.**
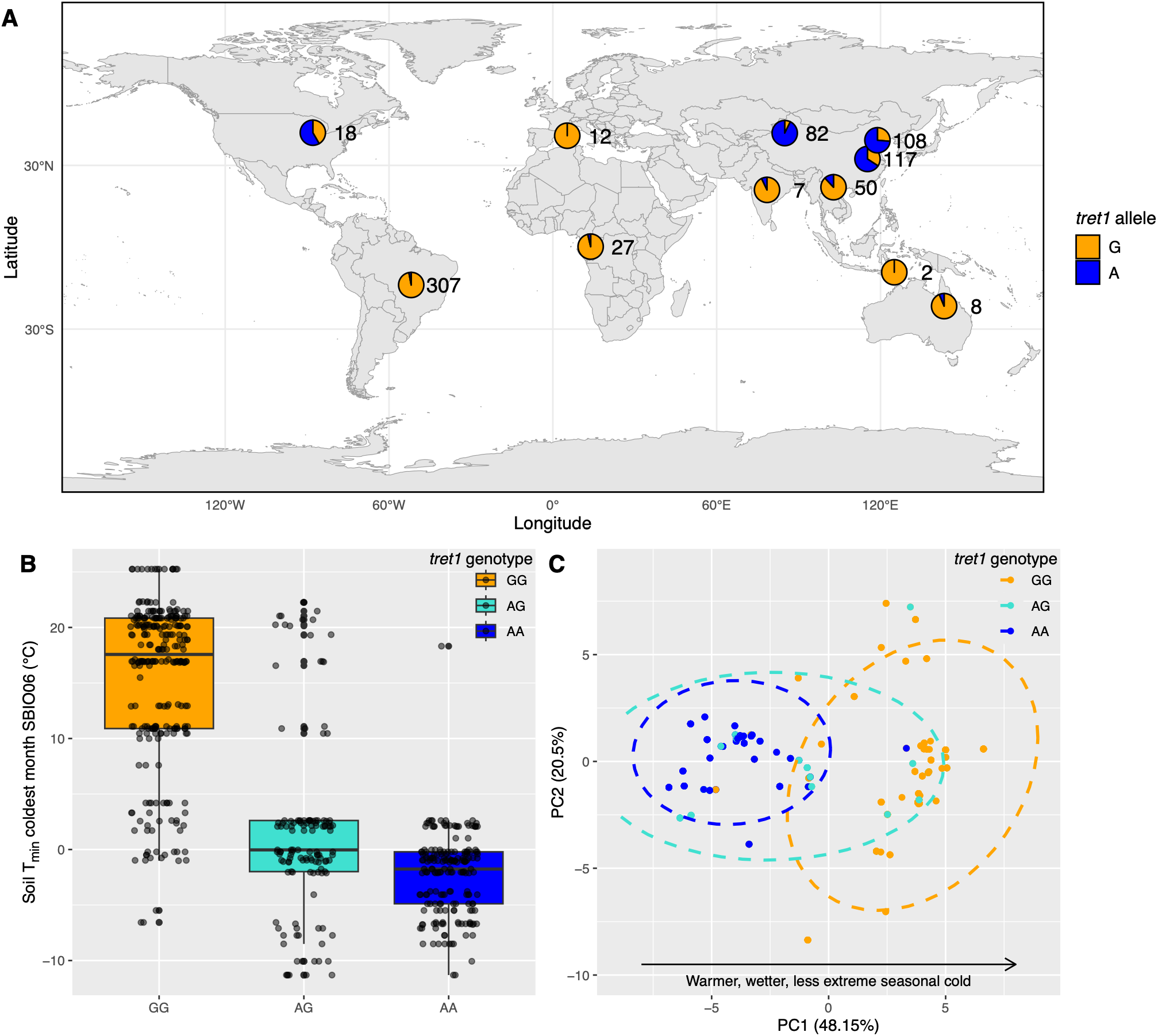
Niche differentiation between wild-type and cold-adapted *H. armigera*. *(A)* Geographic variation in *tret1* allele frequencies (position 3:6626270 in the reference assembly). Sample size from each region is represented by a number next to each pie chart. Individual sampling locations can be found in Figure S3. *(B)* Minimum soil temperature of the coldest month (SBIO06) at a depth of 0-5cm at sampling locations for all three possible *tret1* genotypes. *(C)* PCA of the sampling locations for all individuals in multidimensional niche space. Dotted lines bound the 99^th^ percentile of a multivariate t-distribution fitted to each genotype.

In the introduced range of *H. armigera*, cold-adapted *tret1* is present at low frequency in South America (2.9%, N=307). However, it is much more common in the introduced population at ORD (58.3%, N=18). This differentiation is striking given that South America is the likely source population for the introduction at ORD (24) and cannot be explained by a population bottleneck associated with introduction alone (Fisher’s exact test, p < 10^-15^). We note that this test does not consider heterogeneity in the frequency of this allele in South America, and that there has been little sampling in southern regions such as Argentina where winter minimum temperatures are lower. We also note that we cannot rule out multiple introductions at ORD. Nonetheless, the cold-adapted *tret*-1 allele occurs at low frequencies in all known potential source populations, except for a relatively isolated region of north-western China. Samples from north-western China are genetically distinct and do not cluster with samples from ORD; it is not a plausible source population (24). Therefore, the most parsimonious explanation for the enrichment of cold-adapted *tret1* alleles among individuals at ORD is selection. Consistent with this explanation, the *tret1* locus is an outlier in genome-wide scans of F_ST_ between South America and ORD (Fig S1). Together, these results point to independent local adaptation of *tret1* in the Western and Eastern hemispheres — in invasive and native populations, respectively.

### Niche differentiation between wild-type and cold-adapted *H. armigera*

Because *H. armigera* overwinters as pupae at a shallow depth in soil (26), the most relevant environmental predictor for its year-round persistence is the minimum annual soil temperature. Homozygotes for cold-adapted *tret1* were found at locations with significantly colder minimum soil temperature of the coldest month (SBIO06) than homozygotes for WT *tret1* (Fig. 1B; Mann-Whitney-U test, p < 10^-15^). We also assessed differentiation between carriers of cold-adapted and WT *tret1* in multivariate environmental niche space using PCA and found low niche overlap (Fig. 1C), with differentiation primarily along the first PC. PCA loadings revealed that this PC was strongly associated with environmental predictor variables related to variation in annual temperature (Fig S2). This strong ecological differentiation between alternate *tret1* homozygotes justifies explicitly including *tret1* genotype information in SDM-based predictions of local establishment risk.

### Cold-adapted *tret1* expands the potential range of *H. armigera* in North America

To quantify the contribution of cold-adapted *tret1* to the potential range of *H. armigera* in North America, we used a two-layered sequential SDM approach (29) (Fig. 2). Briefly, this involved training an initial SDM on global occurrences for *H. armigera* to capture the general ecological requirements of the species. Predicted suitability from the initial SDM was then used as a predictor variable in an SDM trained on occurrences of homozygotes for either cold-adapted or WT *tret1*, which captures the specific ecological requirements of each homozygous *tret1* genotype. To differentiate the effects of local adaptation from niche truncation due to restricted geographic sampling, we also trained an SDM on all genotyped occurrences. We did not include *tret1* heterozygotes in our training data for either of the genotypically informed models to avoid making assumptions about the dominance of either *tret1* allele. We fit each SDM using an ensemble modeling approach. Performance metrics (Boyce Index and ROC AUC), and partial dependence curves for each model can be found in Figs. S4-7.

**Figure 2.**
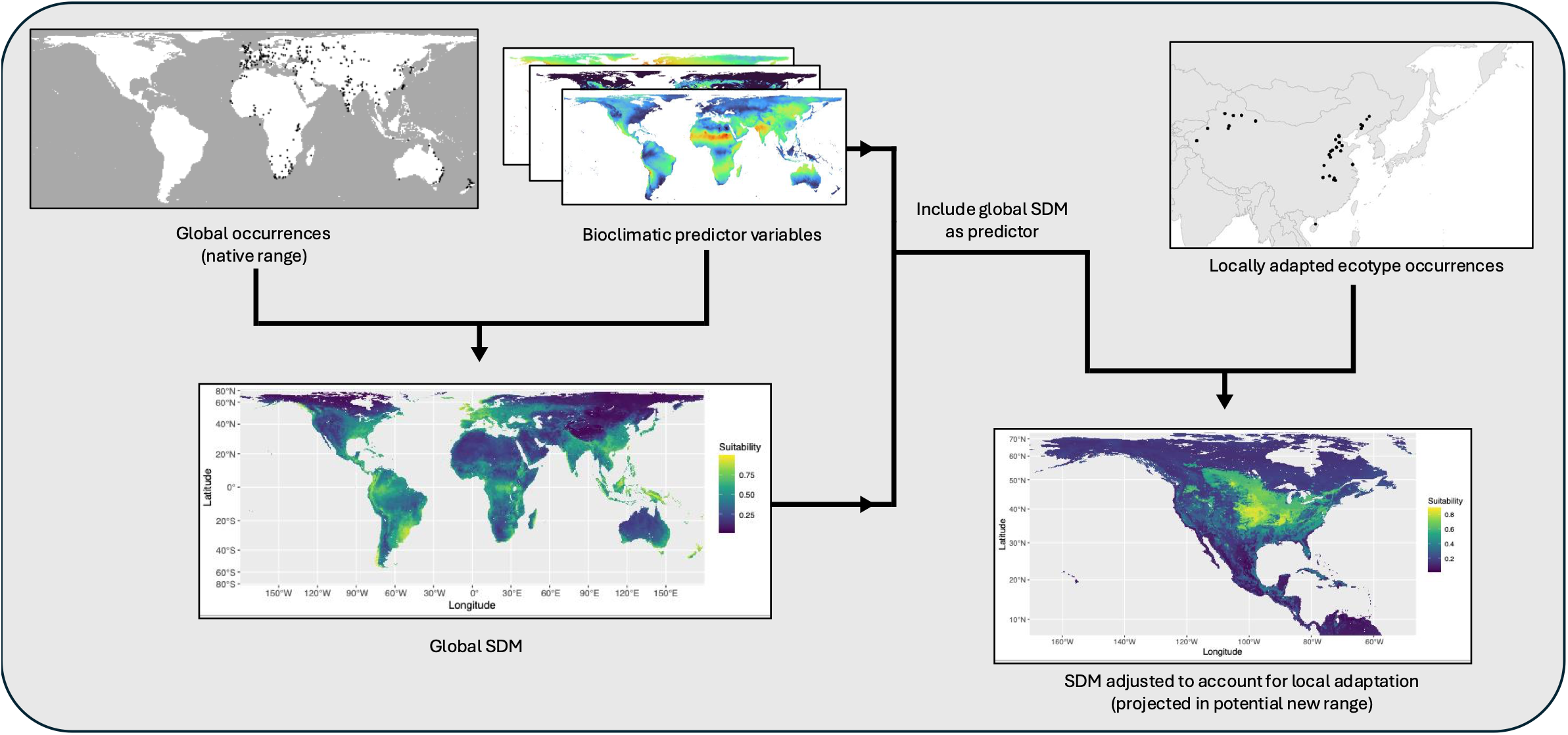
A sequential modeling approach to account for local adaptation in predictions of invasion risk. Using the sequential modeling approach outlined by Guisan *et al*. (2025) (24) overcomes niche truncation arising from restricted geographic sampling of genotyped individuals. An initial SDM is trained on a large set of occurrences for *H. armigera* in its native range, capturing the broad ecological requirements for the species. The prediction from the initial SDM is used as a predictor in a second SDM trained on occurrences carrying a given *tret1* genotype.

Cold-adapted *tret1* dramatically expands the potential range of *H. armigera* in North America (Fig. 3A), doubling the median projected suitability within 100km of ORD as compared to WT *tret1* (Fig. 3B). The two genetically informed models partition the predicted suitability from the uninformed model. WT *H. armigera* has a predicted range primarily in Central America, the Caribbean, and Florida, while cold-adapted *H. armigera* has predicted suitability across the midwestern United States and southern Canada. Notably, cold-adapted *tret1* increases the suitability of the area surrounding ORD, potentially explaining the long-term persistence of the introduced population despite extreme winter temperatures.

**Figure 3.**
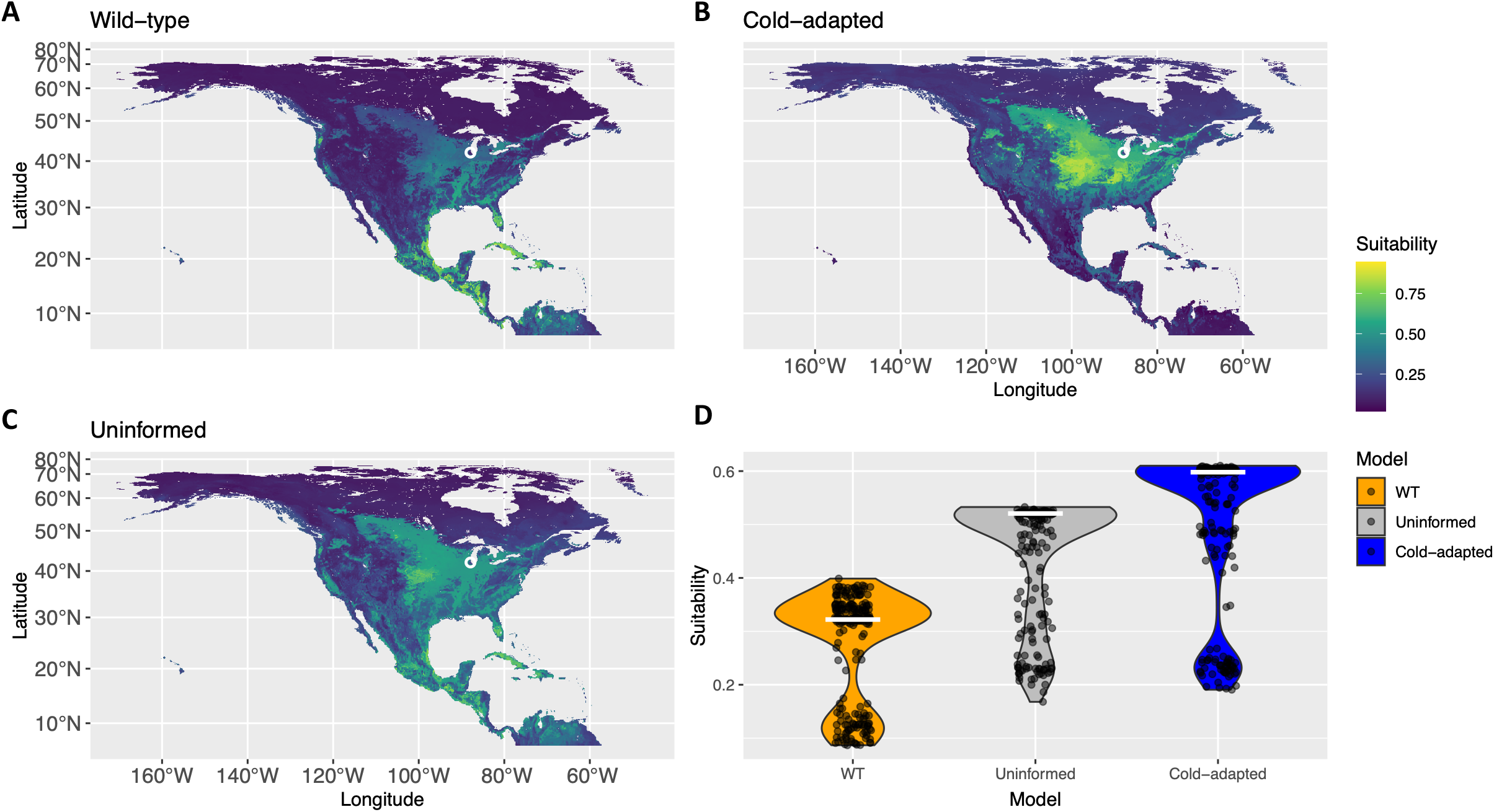
Predicted habitat suitability for wild-type *(A)* and cold-adapted *(B) H. armigera* in present-day North America, and a prediction from an uninformed model that does not consider local adaptation *(C)*. Predictions are the median output of all models in each ensemble with Boyce Index > 0.7. The location of the introduced North American population (ORD) is circled in white. *(D)* Habitat suitability within 100km of ORD for each model. White bars indicate the median of each distribution.

Partial dependence curves for each of the three models show that the main trade-off in suitability between cold-adapted and WT *H. armigera* is driven by opposite responses to minimum soil temperature of the coldest month (SBIO06; Fig. 4), consistent with our assessment of niche differentiation and previous measurements of cold tolerance between the two genotypes (26).

**Figure 4.**
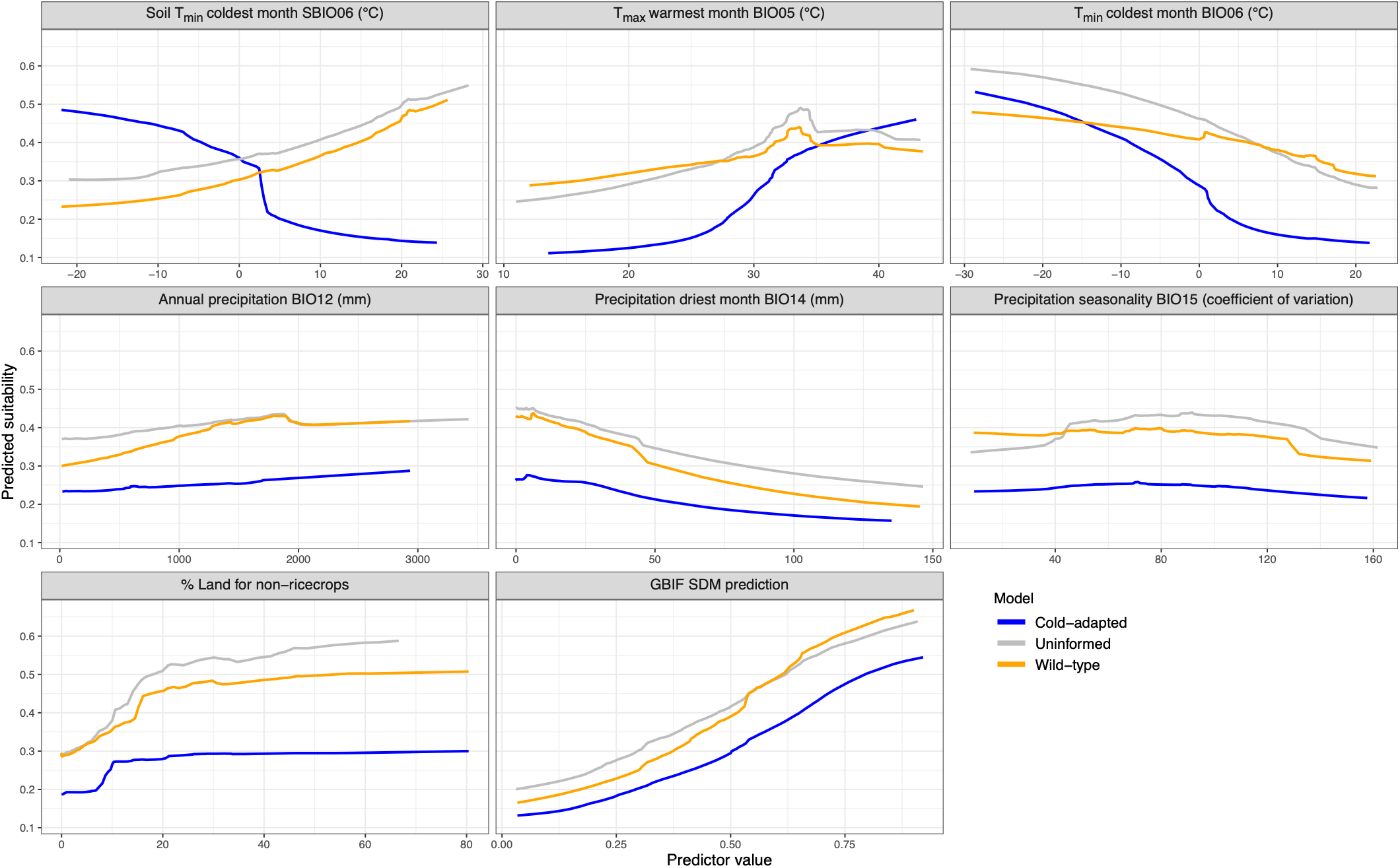
Partial dependence plots showing the predicted marginal habitat suitability response to changing parameters from each model. Marginal responses were calculated by averaging over the background distribution for all other variables from 100 randomly sampled points.

### Cold-adapted *tret1* increases the projected suitable range of *H. armigera* in North America under predicted climate scenarios

To investigate the contributions of WT and cold-adapted *tret1* in *H. armigera* to potential range in North America under future climate scenarios, we retrained each SDM excluding SBIO06 (minimum soil temperature of the coldest month), as there are no future projections available for this predictor. We note that much of the variation in suitability attributed to SBIO06 in the original model is absorbed by BIO06 (the corresponding surface temperature variable; Fig. S6), although SBIO06 and BIO06 were not strongly correlated enough for one of them to be removed to mitigate multicollinearity among predictors. We generated predictions at intervals of 20 years from 2030-2090, across 4 IPCC Shared Socioeconomic Pathways (SSPs; 126, 245, 370, and 585, in increasing order of emissions severity). Continuous predictions of habitat suitability were converted to binary presence/absence predictions by selecting the threshold that maximized the models’ true skill statistic.

While cold-adapted *H. armigera* are predicted to undergo moderate range expansion between 2030 and 2090 under the two most extreme climate scenarios SSP370 and SSP585, there is no significant change in the size of the WT range under any scenario (Fig. 5A). To identify the climate variables driving expansion (or lack thereof) under these scenarios, we computed Shapley values for each predictor variable from 100 randomly selected cells that were either lost, retained, or added to the predicted range. This was repeated for cold-adapted and WT *H. armigera* between 2030 and 2090, across all SSPs. Increases in the maximum temperature of the warmest month (BIO05) and minimum temperature of the coldest month (BIO06) were the main drivers of range gain and loss respectively in cold-adapted *H. armigera* (Fig. 5B-C). Examination of these variables across the sampled cells revealed areas that were lost from the predicted distribution when they became too warm, while those regions that were gained had previously been too cold (Fig. 5C). Even in cells that were predicted to remain suitable, BIO05 took on increasing importance as the main driver of increased suitability. In contrast, WT *H. armigera* seems less sensitive to changes in extreme annual temperature fluctuations, which may explain the stability of its projected range. Overall, these results are consistent with the status of cold-adapted *H. armigera* as having greater fitness in habitats experiencing extreme annual temperature fluctuations.

**Figure 5.**
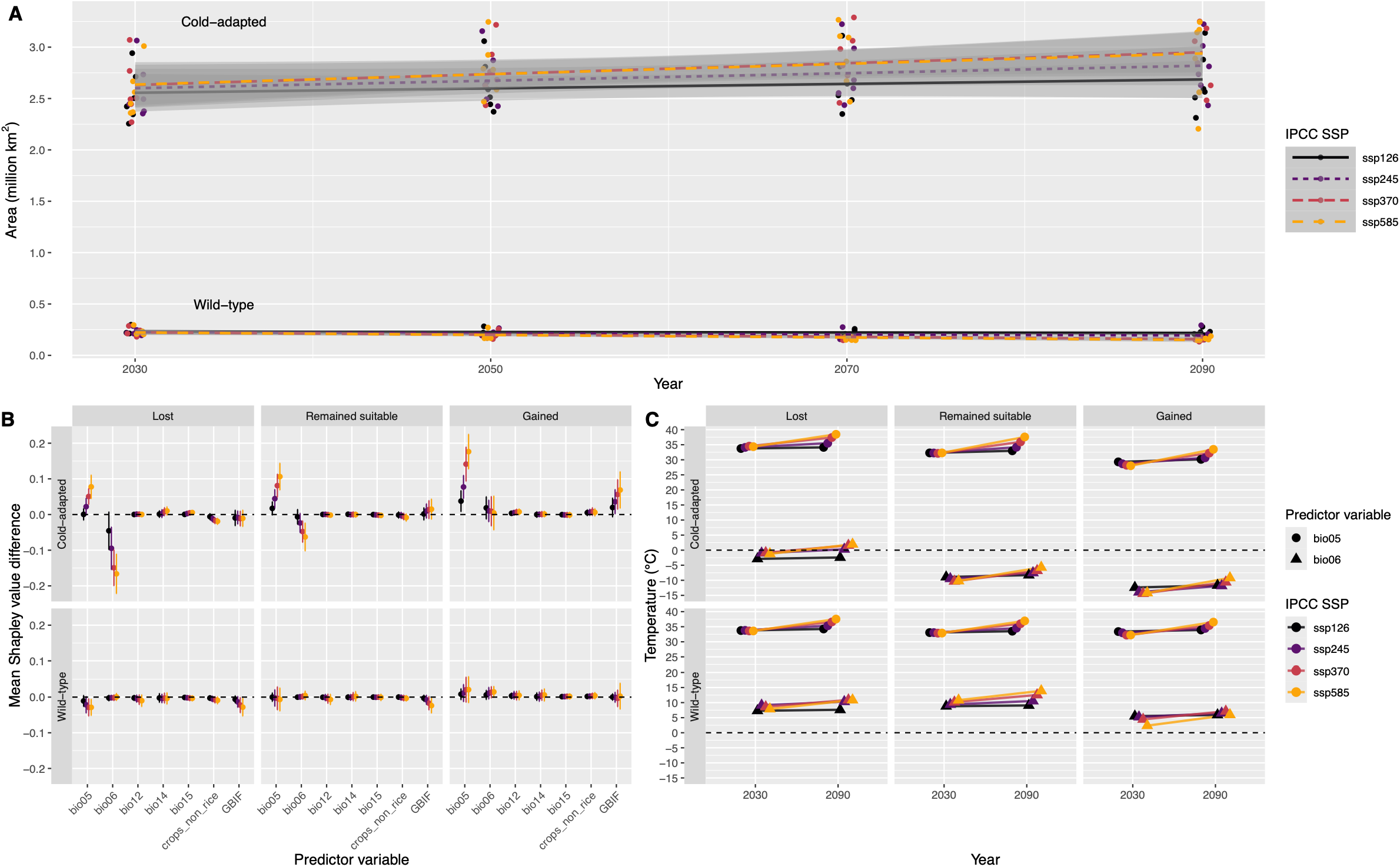
Warming temperatures drive range expansion for cold-adapted but not wild-type *H. armigera*. (A) Predicted suitable area for cold-adapted and wild-type *H. armigera* in North America from 6 CMIP6 models across 4 timepoints and 4 IPCC shared socioeconomic pathways (SSPs), ordered from best-(SSP126) to worst-case (SSP585) scenarios. Continuous predictions of habitat suitability were converted to presence/absence by selecting the threshold that maximized the True Skill Statistic. *(B)* Mean difference in Shapley value for cells that were predicted to be lost, remained suitable, or gained between 2030 and 2090 for cold-adapted and wild-type *H. armigera* for each SSP. Shapley values were calculated as the average difference in prediction between 25 randomly selected coalitions of predictors with and without the focal predictor variable, for 100 randomly selected cells for each scenario. Error bars represent standard deviation. *(C)* Mean change in the two main drivers of range loss or gain (BIO05, maximum temperature of warmest month and BIO06, minimum temperature of coldest month) between 2030 and 2090 for each scenario, calculated from the same cells sampled in *(B)*.

## Discussion

### The role of *tret1* in the persistence of an *H. armigera* population in North America

A major aim in invasion genetics is to understand how evolutionary dynamics contribute to the transport, establishment, and spread of invasive species (30). Here we have developed a novel approach to incorporate known patterns of local adaptation into species distribution models, taking advantage of over 700 whole genome sequences and experimentally established information on a locally adapted allele in *H. armigera*. Leveraging a large whole-genome resequencing dataset, we found that the small *H. armigera* population, currently isolated at ORD, carries the cold-adapted *tret1* allele at a frequency of 58.3%. The frequency of the cold-adapted *tret1* allele is high at ORD compared to its low frequency in South America, which appears to be at least one of the source populations (24). It is therefore likely that there has been positive selection on the cold-adapted allele in the cold winter conditions at ORD, though larger genomic sample sizes in future work are required to verify this conclusion. The current data show that the spatial sorting of alleles has resulted in parallel geographic clines in the Western and Eastern hemispheres, in invasive and native populations, respectively. Consistent with this observation, our suitability models predict parallel clines in both Asia and North America, with decreasing suitability for the cold-adapted *tret1* allele below a latitude of ~30°N. We also note, however, that winter conditions were relatively mild in the region surrounding ORD in the period 2019-2023, though winter temperatures were more extreme in 2025-26 (31). We predict that this recent extreme minimum winter temperature will increase the frequency of the cold-adapted allele.

An interesting avenue for future research would be to test whether other loci show similar repeated clines in the native and invasive populations. We have shown that standing genetic variation was first exported from a cold-adapted native-range source population and maintained at low frequency in invasive South American bridgehead populations. These variants can potentially be ‘re-assembled’ by natural selection into co-adapted allelic combinations that increase fitness in the non-native high-latitude population at ORD — a phenomenon that Schluter and Conte termed the ‘transporter process’ (37). We hypothesize polygenic adaptation to high-latitude environments will proceed via this mechanism.

Although the cold-adapted genotype clearly enhances fitness at higher latitudes through its effect on the supercooling point of hemolymph, we do not have direct evidence for a fitness trade-off in warmer conditions. Regardless of whether the marginal fitness of WT genotype is enhanced or neutral in warmer regions, the expansion of the ecological niche of *H. armigera* attributable to cold-adapted *tret1* is in turn responsible for a dramatic northward expansion of its potential geographic range in North America. The presence of the cold-adapted allele in the founding population doubles the projected suitability of the greater Chicago area, compared to a counterfactual scenario where only WT individuals were introduced. The projected suitability can be interpreted as a proxy for the likelihood that a viable founding population persists. Therefore, the presence of the cold-adapted *tret1* allele has contributed to the overwinter survival and persistence of *H. armigera* at ORD. This finding provides insight into the evolutionary dynamics that govern which introduced populations pass the “filter” from transport to survival in a new range.

In addition to its role in aiding persistence, our suitability models also imply a potential role for *tret1* in facilitating the transport of *H. armigera*. The predicted distribution for WT *H. armigera* suggests the South American population could expand throughout Central America, the Caribbean, and Florida. If the expanding population continues to carry cold-adapted *tret1* at a low-frequency (as has been the case in Brazil), selection for cold-adaptation at the edge of this expanded range could increase its frequency and facilitate spread as far north as Canada. In this hypothetical scenario, selection on different alleles at the same locus would facilitate transport and establishment, respectively. This highlights the importance of understanding the genetic architecture of local adaptation when studying the evolutionary dynamics underlying biological invasion.

### Adaptation at range margins: implications for invasive species

A wealth of theoretical and empirical evidence supports the idea that populations at species range margins are likely to be adapted to extreme climatic conditions, and that range expansion under climate change may be primarily driven by these margin-adapted individuals (38–41). We find this to be the case in *H. armigera*, with range expansion under potential future climate change likely to be driven by cold-adapted individuals at the range margin. Much research in conservation science has focused on the need to preserve these locally adapted range-margin populations, to facilitate species range shifts with climate change (41, 42). The inverse of this argument applies for pest and invasive species control strategies: we suggest that, should *H. armigera* spread outside of ORD to other locations, control efforts could be focused on cold-adapted, range-margin populations to inhibit the impacts of any future climate-driven range expansion.

In contrast to the expanding cold-adapted range, WT *H. armigera* in the core of the species’ potential range do not undergo significant expansion under the tested set of future climate scenarios. Such “leading edge” range expansion could result from competition between the cold-adapted and WT individuals, where climate change in the core of the species range is not significant enough to give WT individuals an advantage over their cold-adapted competitors (41). Failure of the WT range to expand could also be due to sensitivity to extreme climatic events which are predicted to increase in frequency if increasing global temperature tends continue (42, 43).

### Sequential SDMs as an evidence-based framework for genetically informed prediction of invasive spread

The ease with which large landscape-genomic datasets can now be generated has led to increasing interest in incorporating information about local adaptation into predictions of species responses to climate change. The sequential modeling approach developed here overcomes limitations associated with previous approaches including niche truncation and lack of interpretability. Furthermore, it limits predictions to loci that have been causally linked to local adaptation, thereby avoiding false signals that result from population structure alone. Our approach integrates information from the distributions of genotyped samples as well as the whole species, explicitly incorporating prior information about local adaptation, and providing an interpretable result about the distributions of carriers of — in this case — the two *tret1* alleles.

A limitation of our approach is its consideration of adaptation at a single locus, while adaptation to climate can be a highly polygenic quantitative trait (44). However, in at least some cases a large proportion of genetic variance locally adapted such can be explained by a relatively small number of large-effect loci — as is the case in this species for low-temperature survival (32) — so our approach provides a useful first approximation. Future developments in genetically informed SDMs would benefit from considering multiple loci that are known to underlie local adaptation (45). One approach could be to group samples by their genotypes at multiple loci, however as the number of loci under consideration increases, the number of groups grows exponentially and so the number of samples per group will become very small. An alternative could be a multi-layered sequential SDM approach, with an additional layer for each locus under consideration. How such an approach would perform with many loci, each with potentially epistatic interactions is unclear, though this could be tested using simulation approaches.

Despite these limitations, this work represents a useful step towards integrating genomic and geospatial data to produce interpretable predictions about the potential spread of an invasive crop pest. We show that considering intraspecific variation in responses to climatic variables can substantially change predictions of suitable habitat. We advocate that future risk assessments and management strategies proactively consider the role of evolutionary processes in shaping the distribution of problematic species.

## Materials and Methods

### *tret1* SNP genotyping

We analyzed short-read whole-genome resequencing data from *H. armigera* individuals reported in North *et al*. (2024) (30), Jin *et al*. (2023) (32), Anderson *et al*. (2016) (46) and North *et al*. (2024) (45). These samples were collected between 2002 and 2021 from 13 countries (Supplementary table S1). We trimmed raw reads to remove Nextera adapter sequences and poly-G tails from lack of Illumina signal using fastp v1.0.1 (47). We removed reads with > 5 missing base calls, or > 40% PHRED quality < 15, or length < 50bp. We aligned remaining reads to the *H. armigera* reference genome (32) using bwa v0.7.19 (48). We sorted the resulting BAM files using SAMtools v1.22 (49) and removed duplicate reads using Picard v3.4.0 (https://broadinstitute.github.io/picard/) before indexing. We estimated genotypes in a ~35.4kb region on Chr3 containing *tret1* (3:6614896-6649427) simultaneously across all individuals using mpileup and call from bcftools v1.21 (49).

We applied filters of variant and genotype PHRED quality >= 30, variant mean depth between 5x and 15x, and genotype depth between 3x and 20x using VCFtools v0.1.16 (50). We imputed missing genotypes at the focal *tret1* SNP (3:6626270) using BEAGLE v5.5 (51).

### Sequential distribution modeling

#### Climate data

We obtained present-day global bioclimate data (BIO01-BIO19) from the WorldClim database v2.1 (52), and global distributions of irrigated and rain-fed non-rice crops from 2015 from HYDE v3.2 (53) at 10 arc-minute resolution using pastclim v2.2.0 (54). Because *H. armigera* overwinters as pupae in soil, we also obtained present-day projections of minimum annual soil temperature at 0-5cm depth (SBIO06) at 2.5 arc-minute resolution from Lembrechts *et al*. (2022) (55). We reprojected all geographic data to the EPSGG-6933 Equal Earth Projection (56) with 10km resolution to prevent sampling bias due to uneven grid cell sizes.

We associated georeferenced samples with climate data from the appropriate locations using terra v1.8 (57). Anderson *et al*. (2016) only provided location data at the level of municipality or city. Since the per-generation dispersal of *H. armigera* is a similar scale to the width of these geographic regions (10s-100s of km) (27), these location data were still informative; we assigned locations at random within the recorded sampling region. We excluded individuals sampled from South America before 2017 (~30-50 generations post-introduction), because the location of individuals sampled shortly after introduction may be more reflective of the location of introduction, rather than the individuals’ environmental niche, violating the assumption of niche-environment equilibrium.

#### Global occurrence data

We retrieved occurrence records for *H. armigera* in its native range between 2000 and 2026 from the Global Biodiversity Information Facility (GBIF) database in March 2026 (58). We kept only records describing a presence (“OccurenceStatus is Present”), with latitude/longitude reference (“HasCoordinate is true”), without spatial issues (“HasGeospatialIssue is false”), and for which the field “BasisOfRecord” was “Human Observation”. To avoid further potential spatial issues, we used CoordinateCleaner v3.0.1 (59) to exclude coordinates that met any of the following conditions: 10km around country capitals, 1km around country centroids, 100km around GBIF headquarters, 100m around biodiversity institutions, or 1 degree around (0,0). We also excluded occurrences in the sea and those with identical longitude and latitude. We combined this dataset with the locations of our genotyped samples and then thinned all occurrences by 30km to reduce the effect of spatial sampling bias to produce our final global occurrence dataset.

#### Global predictor variable selection

We selected an initial set of candidate predictor variables (SBIO06, BIO06, BIO05, BIO01, BIO12, BIO14, BIO15, crops_non_rice, BIO04; interpretation of variable codes can be found in Supplementary Table S2) *a priori* based on the ecology of *H. armigera* and previous distribution modeling work for this species (33). To reduce issues arising from multicollinearity among predictor variables in correlative models, we calculated Pearson’s correlation coefficient between all pairs of variables among occurrence locations and selected only a single variable within each group defined by a correlation cutoff of 0.7. BIO01 was the only variable removed this way.

#### Global distribution modeling

We fitted a species distribution model to our global occurrence dataset using an ensemble modeling approach implemented in tidysdm v1.0.4 (60). Algorithms included in the ensemble were: regularized regression (RGLM, glmnet v4.1 (61, 62)), generalized additive models (GAM, mgcv v1.9 (63)), random forests (RF, ranger v0.17 (64)), gradient boosting machines (GBM, xgboost v1.7.8 (65)) and maximum entropy (Maxent, maxnet v0.1.4 (66)). We selected these algorithms to provide a good balance between regression and classification approaches. We sampled calibration data and fitted the models using the following steps:

##### 1: Thinning occurrences

Although spatial thinning can reduce the impacts of spatial sampling bias, there is no way to determine the optimal thinning distance (67), and previous work has shown it is often beneficial to additionally filter in environmental as well a geographical space (68). Therefore, we thinned occurrences in multivariate niche space by conducting PCA on the selected predictor variables at occurrence locations, and randomly selecting one occurrence from each square of a 30×30 grid projected onto PC1 and PC2.

##### 2: Background sampling

We randomly sampled background points from the unioned buffer of 500km (selected based on the plausible per-generation dispersal distance for *H. armigera* (27)) radii around each occurrence. We sampled 10× as many background points as occurrences. As all the algorithms we used except Maxent can be sensitive to class imbalance, we balanced classes by down-weighting the background points for the RGLM, GAM, and GBM, and down-sampling them for the RF. We did not allow Maxent to use hinge transformations of predictor variables as they are prone to overfitting.

##### 3: Cross-validation

We tuned each model in the ensemble using 5-fold spatially blocked cross-validation on a 10×10 spatial grid. 10 combinations of hyperparameters were explored using a grid search for each model with the goal of maximizing model performance, measured by the Boyce Index (69), which we selected as the best-performing metric in the context of presence-only SDMs (70). We recorded both Boyce Index and ROC AUC for each calibrated model.

##### 4: Bootstrapping

Because SDMs can be sensitive to stochasticity introduced by the effect of environmental thinning (sampling bias in species occurrence records) and background sampling, we performed 10 replicates of steps 1-3. From each replicate, only models with mean Boyce Index > 0.7 across all folds were included in the final ensemble. Final consensus projections were taken as the median projection of all included models.

#### Genotype-informed distribution modeling

We fitted two separate distribution models to occurrences for individuals homozygous for alternate *tret1* alleles. Additionally, we fitted a third distribution model to all genotyped occurrences, to differentiate between the effects of local adaptation and limited geographic sampling. The distribution modeling approach was identical to that for the global SDM, with the following exceptions:

1. We used the output of the global SDM as a predictor variable. We therefore carried out another round of removing correlated variables, resulting in the removal of BIO04.
2. We did not spatially thin occurrences as there are relatively few (homozygous WT N=58, homozygous cold-adapted N=38) compared to the global SDM and instead relied on environmental thinning alone.
3. In locations with many occurrences for one genotype and few or none from the other, the absence of one genotype is due to biological reality rather than lack of sampling effort (i.e. these locations carry information about climatic requirements for the absent genotype). Therefore, for each genotypic SDM, we also sampled background points from the unioned buffer around occurrences for the opposite genotype. To prevent the generation of folds with no occurrences, we used random rather than spatially blocked 5-fold cross-validation.

To examine the contributions of individual variables for the global and genotypic SDMs, we constructed partial dependence curves using DALEX v2.5.3 (71). Marginal responses were calculated by averaging over the background distribution for all other variables from 100 randomly sampled points.

#### Projection into future climates

Because future projections for soil bioclimate variables are not available, we retrained all models with SBIO06 removed as a predictor variable.

We obtained future bioclimate data (BIO01-BIO19) projections from the WorldClim database v2.1 (52) for 4 time periods (2021-2040, 2041-2060, 2061-2080, 2081-2100) and 4 IPCC shared socioeconomic pathways (SSPs; 126, 245, 370, 585, ordered from best-to worst-case climate scenarios) (43). We obtained these data for 6 different climate projections from the Coupled Model Intercomparison Project Phase 6 (CMIP6) (72), selected based on recommended best practice for North America (73): EC-Earth3-Veg (74), GISS-E2-1-G (75), ACCESS-CM2 (76), MRI-ESM2-0 (77), MPI-ESM1-2-HR (78) and MIROC6 (79). We also used the latest available crop distribution data from HYDE v3.2 (53), which is from 2023.

We created projections of future habitat suitability across all time periods, SSPs, and climate projections for the two retrained genotypic SDMs. Consensus projections for each time and SSP were created by taking the median across all underlying climate projections. Projections were also converted to presence-absence by selecting the projected suitability threshold that maximizes the true skill statistic (TSS).

#### Identifying drivers of range expansion under climate change

To identify which predictor variables drive predicted range changes, we calculated Shapley values (80) for each predictor using DALEX v2.5.3 (71) for cells that were predicted to be either lost, remain suitable, or gained between 2030 and 2090 for each combination of CMIP6 projection, SSP and wild-type or cold-adapted *H. armigera* (=3 x 6 x 4 x 2 = 144 scenarios). We computed Shapley values as the mean difference in predicted suitability across 25 random coalitions of predictor variables when the focal variable was included or excluded, for 100 cells from each scenario.

## Acknowledgments

We thank Todd Gilligan, Luke Tembrock, Frida Zink, and Brad Coates for their extensive support and detailed feedback on this project. We also thank Andrea Manica, Michela Leonardi, Arman Pili, and Nik Cunniffe for their feedback and advice on the analysis. The findings and conclusions in this publication are those of the authors and should not be construed to represent any official USDA or U.S. Government determination or policy.

## Code availability

All scripts are available at https://github.com/cd-williams/helicoverpa-tret1-sdm

## Notes

### Competing Interest Statement

The authors have declared no competing interest.

