## Supplementary materials for "Genetically informed distribution models refine predictions for the overwintering range of *Helicoverpa armigera* in North America"

### Supplementary figures

#### Fig S1:

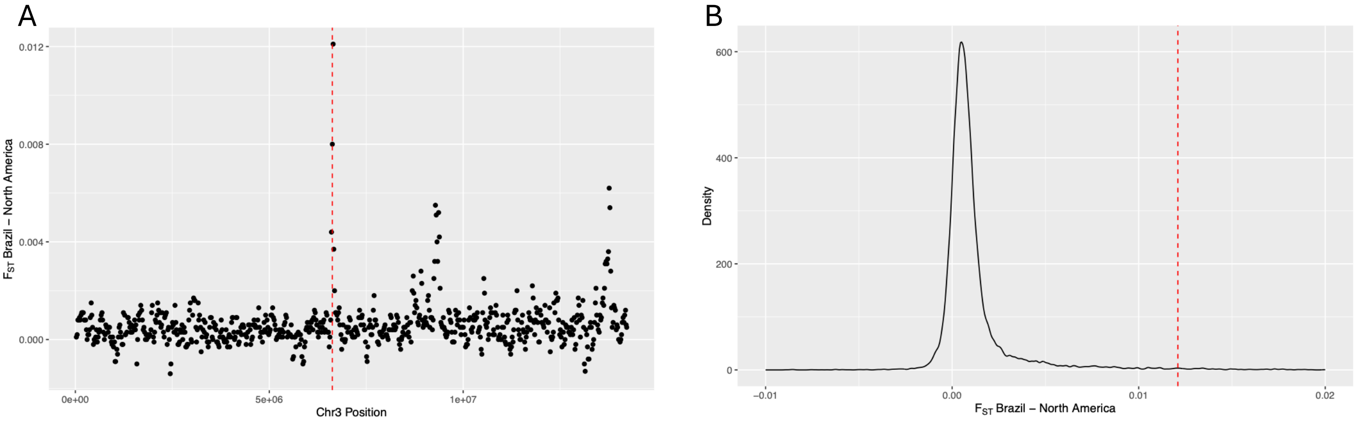


**Figure S1: Differentiation at the *tret1* locus between Brazil and North America.** *(A)* $F_{ST}$ between Brazilian and North American populations in a sliding window (size=50kb, step=20kb) along Chr3. *tret1*is marked with a dashed red line. *(B)* Genome-wide distribution of $F_{ST}$ between Brazilian and North American populations (sex chromosomes excluded). The maximum value of the *tret1* peak from *(A)* is marked with a dashed red line.

#### Fig S2:

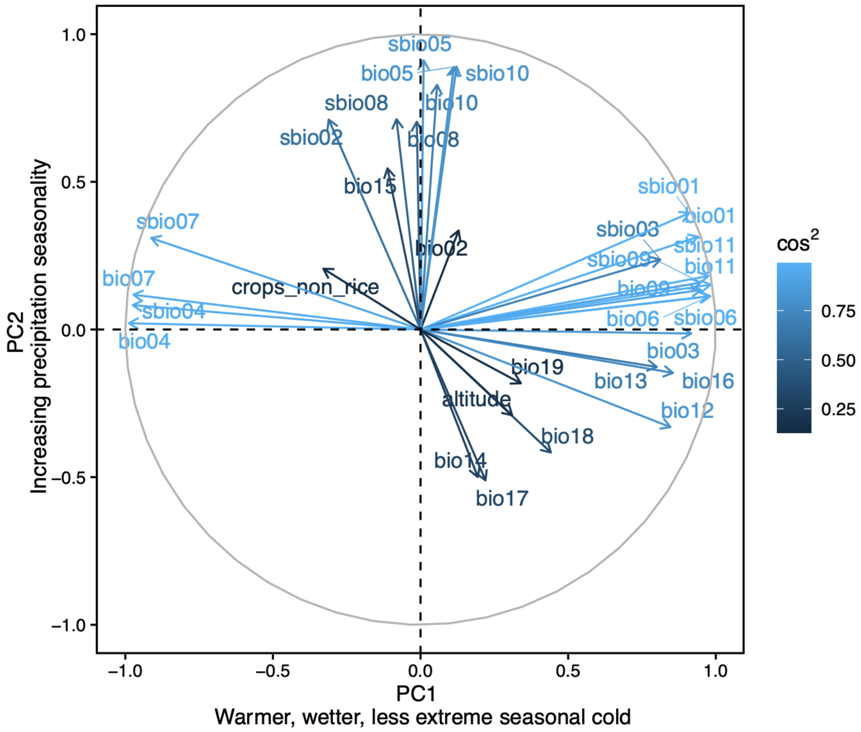


**Figure S2:** Loadings from the PCA in Fig. 1C. Variable codes are matched to variable names in Table S2.

**Fig S3:**

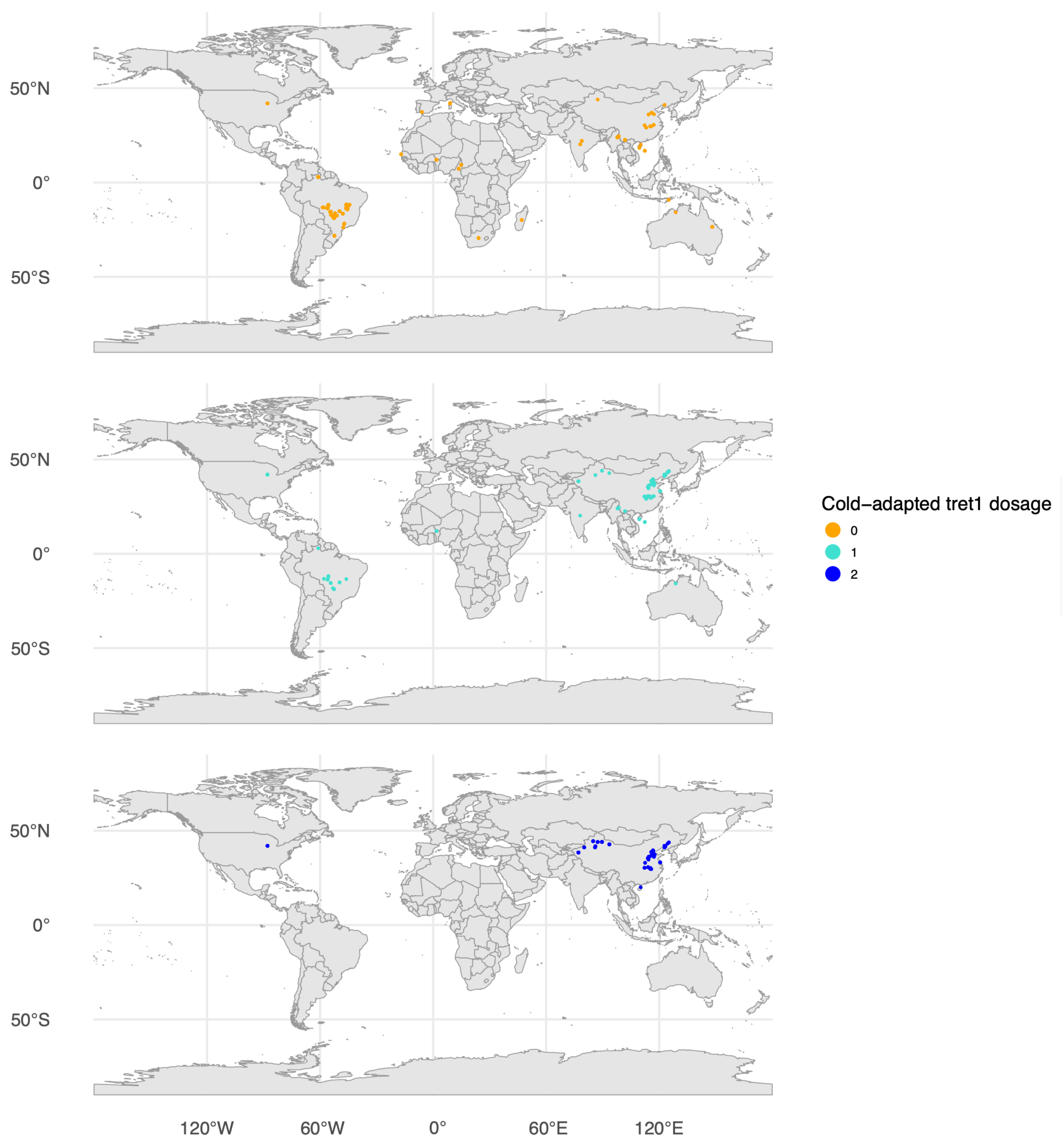


**Figure S3:** Locations of individuals carrying each of the 3 possible *tret1* genotypes

### Fig. S4

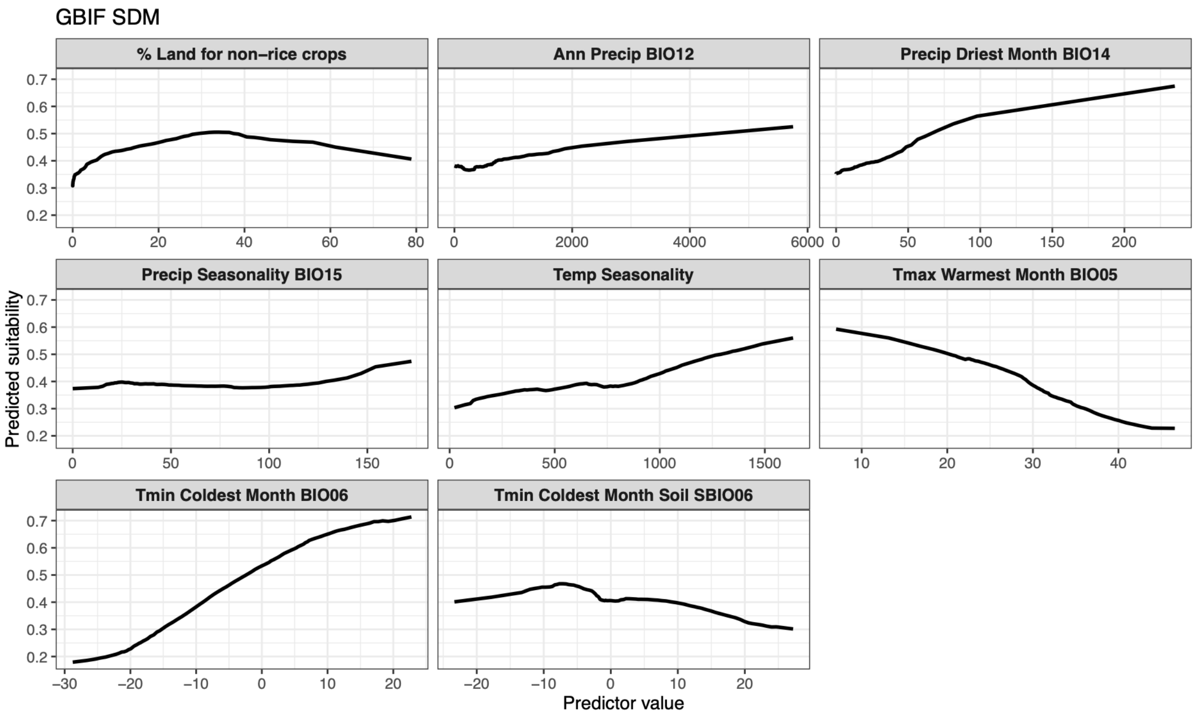


**Figure S4:** Predicted marginal responses to changing climatic conditions for the GBIF SDM. Calculated the same way as Figure 4.

### Fig. S5

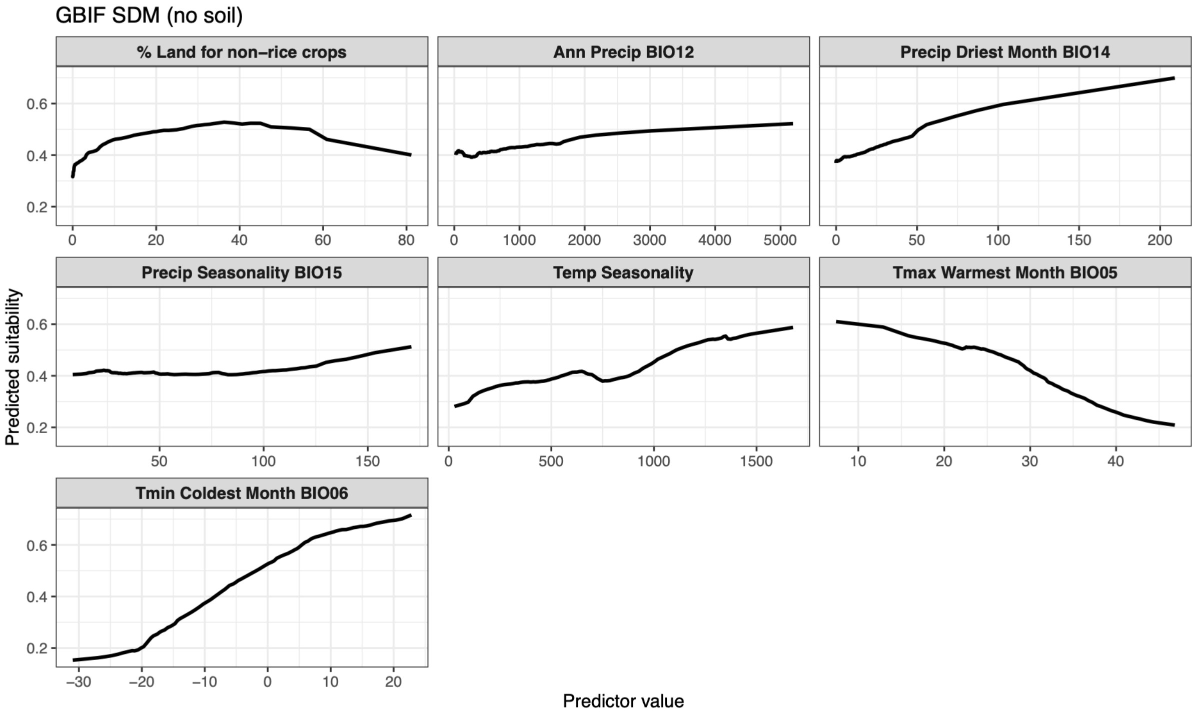


**Figure S5:** Predicted marginal responses to changing climatic conditions for the GBIF SDM, with soil temperature variables excluded. Calculated the same way as Figure 4.

### Fig. S6

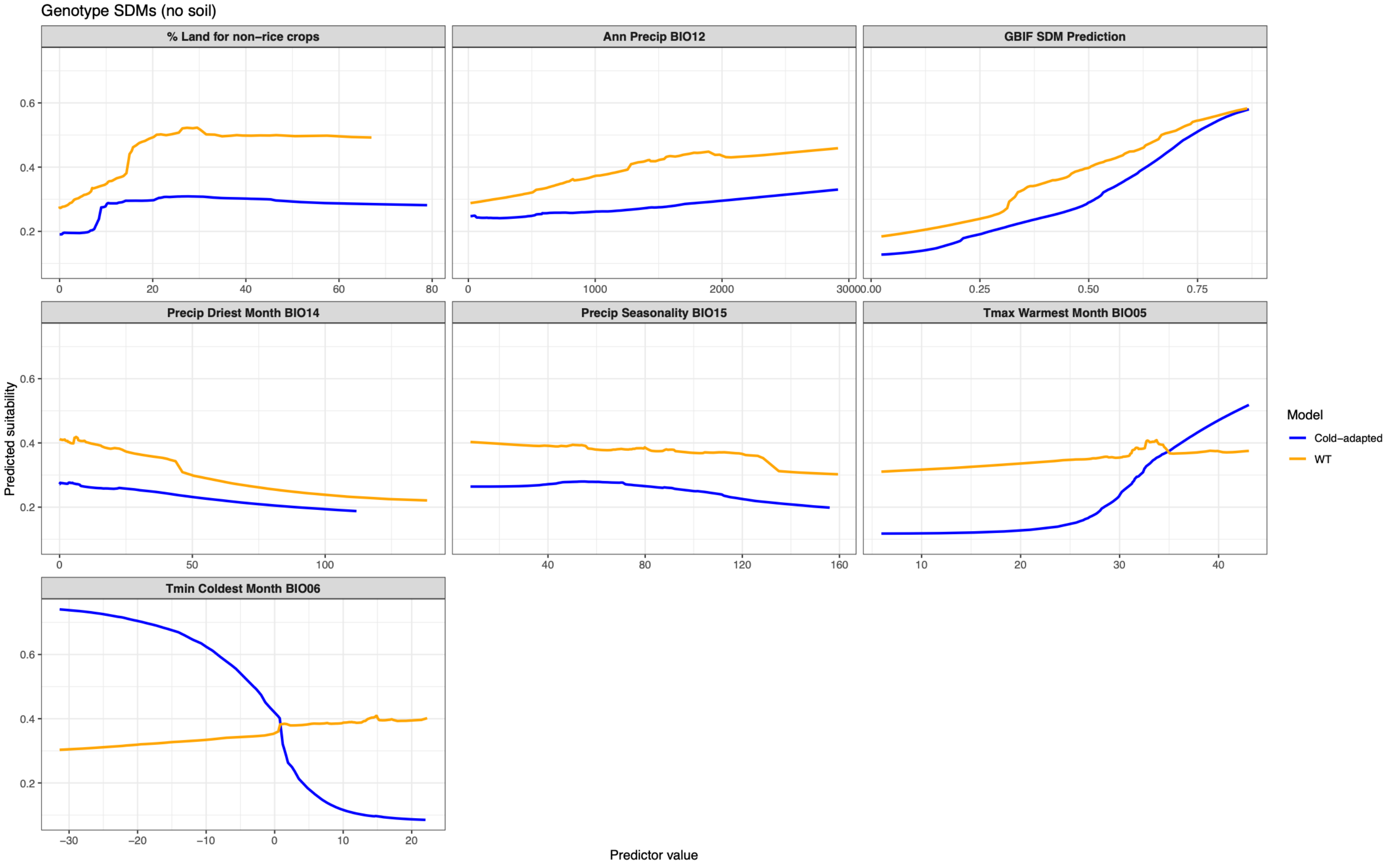


**Figure S6:** Predicted marginal responses to changing climatic conditions for the each of the genetically informed SDMs, with soil temperature variables excluded. Calculated the same way as Figure 4.

### Fig. S7

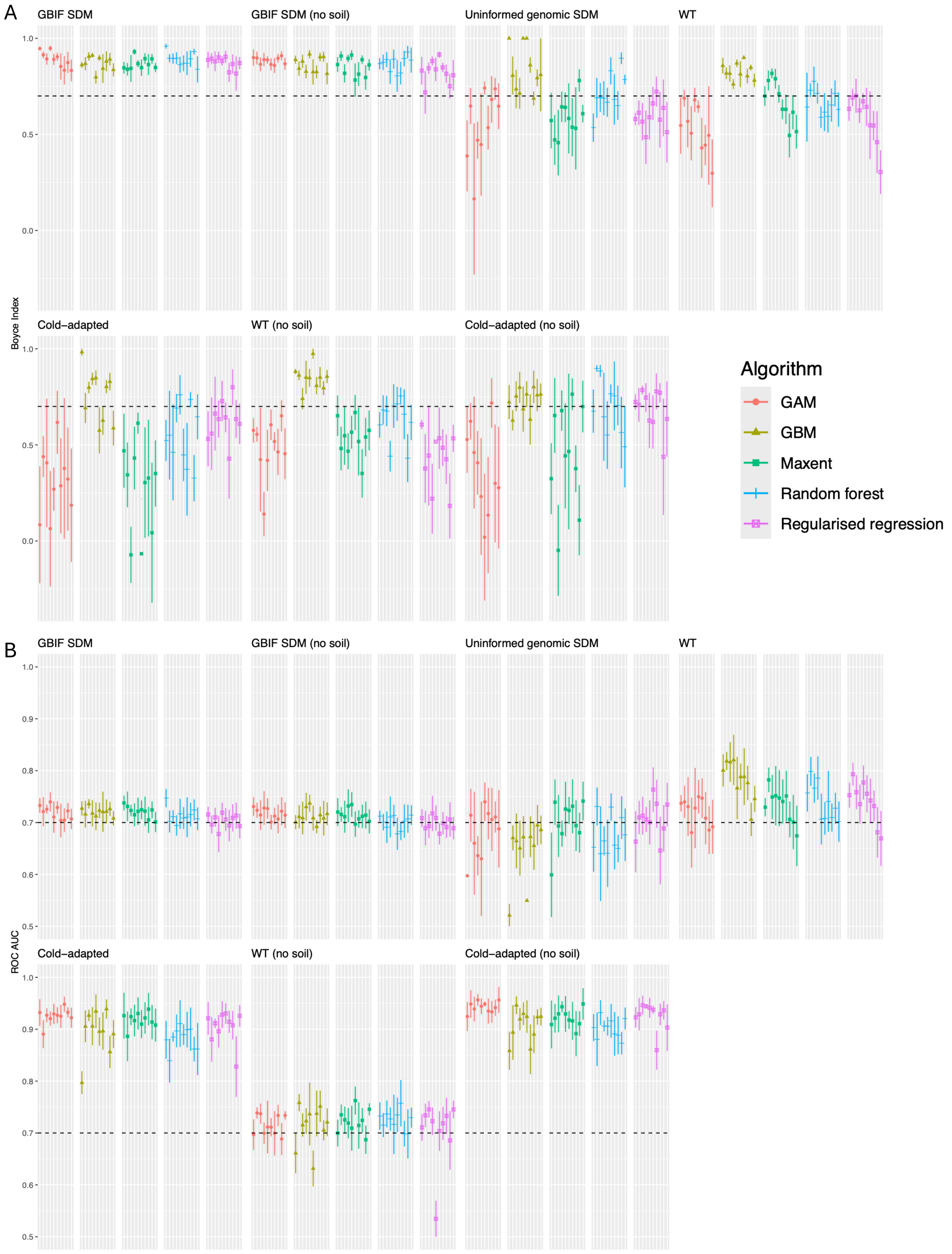


**Figure S7:** Performance of all models measured by (A) Boyce index and (B) ROC AUC. Only models with Boyce Index > 0.7 (indicated by dashed line) were included in the final ensembles. ROC AUC is not necessarily maximal, since models were fit to maximise the Boyce Index.

### Supplementary tables

**Table S1** see tableS1.csv

**Table S2:** Environmental predictor variable codes and their corresponding variable names. “Soil” variables are at a soil depth of 0-5cm.

| **Variable code** | **Variable name** |
|---|---|
| (s)bio01 | Annual mean (soil) temperature (°C) |
| (s)bio02 | Mean diurnal (soil) temperature range (°C) |
| (s)bio03 | (Soil) isothermality (%) |
| (s)bio04 | (Soil) temperature seasonality *(standard deviation)* |
| (s)bio05 | Maximum (soil) temperature of warmest month (°C) |
| (s)bio06 | Minimum (soil) temperature of coldest month (°C) |
| (s)bio07 | (Soil) temperature annual range (°C) |
| (s)bio08 | Mean (soil) temperature of wettest quarter (°C) |
| (s)bio09 | Mean (soil) temperature of driest quarter (°C) |
| (s)bio10 | Mean (soil) temperature of warmest quarter (°C) |
| (s)bio11 | Mean (soil) temperature of coldest quarter (°C) |
| bio12 | Annual precipitation (mm) |
| bio13 | Precipitation of wettest month (mm) |
| bio14 | Precipitation of driest month (mm) |
| bio15 | Precipitation seasonality *(coefficient of variation)* |
| bio16 | Precipitation of wettest quarter (mm) |
| bio17 | Precipitation of driest quarter (mm) |
| bio18 | Precipitation of warmest quarter (mm) |
| bio19 | Precipitation of coldest quarter (mm) |
| altitude | Altitude (m) |
| crops_non_rice | Proportion of land covered by non-rice crops, both irrigated and rain-fed (%) |
